# Orthogonalization of spontaneous and stimulus-driven activity by hierarchical neocortical areal network in primates

**DOI:** 10.1101/2024.07.01.601463

**Authors:** Teppei Matsui, Takayuki Hashimoto, Tomonari Murakami, Masato Uemura, Kohei Kikuta, Toshiki Kato, Kenichi Ohki

## Abstract

Understanding how biological neural networks perform reliable information processing in the presence of intensive spontaneous activity (*1-3*) is an essential question in biological computation. Stimulus-evoked and spontaneous activities show orthogonal (dissimilar) patterns in the primary visual cortex (V1) of mice (*4-6*), which is likely to be beneficial for separating sensory signals from internally generated noise (*5, 7-12*); however, those in V1 of carnivores and primates show highly similar patterns (*3, 13-17*). Consequently, the mechanism of segregation of stimulus information and internally generated noise in carnivores and primates remain unclear. To address this question, we used primate-optimized functional imaging (*18*) to precisely compare spontaneous and stimulus-evoked activities in multiple areas of the marmoset visual cortical pathway (*19*). In marmoset V1, but not in mouse V1, spontaneous and stimulus-evoked activity shared similar activity patterns, suggesting a function of spontaneous activity specific to mammals with functional columns. However, in the higher-order visual areas of marmosets, spontaneous and stimulus-evoked activities exhibited dissimilar patterns. Analysis of neural activity geometry further revealed progressive orthogonalization of the two types of activities along the cortical hierarchy in marmosets, which reached a level comparable to that of mouse V1. Thus, orthogonalization of spontaneous and stimulus-evoked activity is a general principle of cortical computation, which, in primates, is implemented by the hierarchical areal network.

## Introduction

Spontaneous activity is one of the most characteristic features of the biological brain that distinguishes it from current artificial neural networks. Studies on global spontaneous activity, which is often observed with functional magnetic resonance imaging (fMRI), collectively revealed the precise similarity between spatial patterns of spontaneous and task-evoked brain networks (*1, 14, 20*). Resting-state activity can be used to predict individual task-evoked fMRI activity (*21*). Such similarity provides important clues for understanding how the intrinsic dynamics of neural circuits may gauge cognitive functions (*1, 21-25*). However, at both the cellular– and mesoscales, the presence of similarity between spontaneous and evoked neural activity remains controversial. Recent studies using cellular resolution calcium imaging reported that, in the primary visual cortex (V1) of mice, patterns of spontaneous activity are orthogonal to those of visually evoked activity (*4-6*). Theoretical considerations indicate that the orthogonal relationship separates stimulus-related signals from internally generated noise, which is advantageous for perceptual stability (*5, 7*). The fact that much of the spontaneous activity in the mouse V1 encoded behavioral information (*4, 6*) further supports the idea that the orthogonal relationship is advantageous for separating stimulus-related and behavioral information.

In contrast to these recent studies, several previous studies that used voltage-sensitive dye imaging and electrode-array recordings showed high similarity between the spatial patterns of spontaneous and stimulus-evoked activity (*3, 13, 16, 17*). Notably, seminal works by Grinvald and colleagues reported that, in V1 of cats and macaques, neuronal activity patterns resembling iso-orientation columns spontaneously appeared without visual stimulation (*3, 15*).

The apparent differences between the findings of the two sets of studies may stem from several key differences. First, there is a difference in experimental techniques: mesoscale functional recording of neural population activity in cats and primates versus two-photon calcium imaging at a cellular resolution in mice. Second, a difference in analytic techniques: the former set of studies mostly used simple spatial correlation, whereas the latter set of studies devised a geometric analysis of neural activity space. Third, there is a potential species-related difference. At the level of local circuits, carnivores and primates possess neocortex with functional columns (*26, 27*), whereas rodents possess non-columnar neocortex (*28, 29*). In addition to these differences, the fact that both sets of studies were conducted only on V1 makes it difficult to extend these results to general cortical computations. Most importantly, if spontaneous and evoked activities share similar patterns in all the visual areas of carnivores/primates, it would be of great interest to determine how the cortical network could distinguish between stimulus-related information from internally generated noise.

Therefore, we aimed to distinguish between these possibilities and to gain insights into the principles of cortical computations by investigating the relationship between spontaneous and stimulus-evoked activity in multiple cortical areas comprising the hierarchical visual network of marmoset monkeys. Marmoset monkeys have a visual network with a well-defined hierarchy (*19, 30*). Unlike macaques, the majority of visual areas in marmosets are exposed on the cortical surface and are thus accessible using optical approaches (*18, 31-35*). To systematically characterize the spatiotemporal patterns of spontaneous activity across the marmoset visual areas, we used a newly developed primate-optimized expression system of genetically encoded calcium indicators (Hashimoto et al., in preparation).

## Results

To observe spontaneous and stimulus-evoked activity across the cortical network, we used marmoset monkeys whose flat neocortex allows easy optical access to many cortical areas (*19*). Grid-based injection of adeno-associated virus carrying primate-optimized GCaMP (Tandem-GcaMP (*36*); Hashimoto et al., in preparation) (**Fig. 1a**), a genetically encoded calcium indicator, successfully transduced a large contiguous cortical volume spanning the occipital and parietal lobes (**Fig. 1b**). Wide-field imaging with visual stimulation revealed patchy functional maps in multiple visual areas (**Fig. 1c**), demonstrating columnar-scale functional resolution throughout the large field of view (FOV).

**Figure 1.**
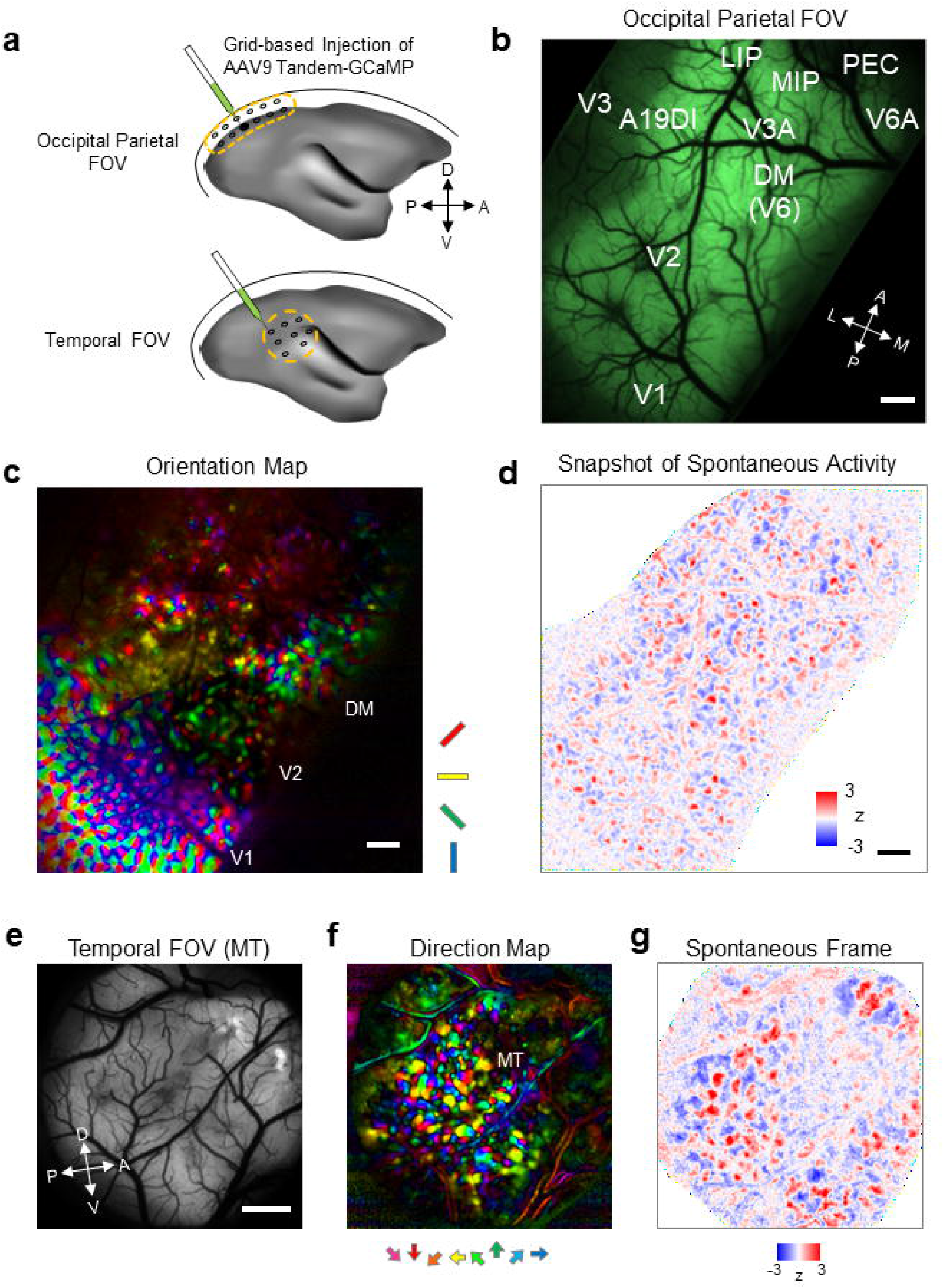
Mesoscale functional imaging revealed patchy spatial patterns of spontaneous activity throughout the marmoset neocortex. **a)** Schematics of large FOV calcium imaging using a primate-optimized GCaMP-expression system. Grid-based injection allowed to cover large surface areas in an occipito-parietal region (top) and a temporal region (bottom). **b)** in vivo GCaMP fluorescence in the occipital parietal FOV. Approximate positions of cortical areas are based on the marmoset brain atlas (*75*). Scale bar, 1 mm. **c)** Orientation map of the occipital parietal FOV obtained with *in vivo* widefield calcium imaging. Positions of V1, V2, and DM are determined based on the structure of the orientation map (note the stripes in V2). Scale bar, 1 mm. **d)** A snapshot of spontaneous activity in the occipital parietal FOV. A spatial high-pass filter was applied. See **Supplementary Video 1** for non-filtered data. Scale bar, 1 mm. **e)** In vivo GCaMP fluorescence in the temporal FOV. Scale bar, 1 mm. **f)** Direction map in the temporal FOV. Direction columns indicate the position of MT. 1 mm. **g)** A snapshot of spontaneous activity in the temporal FOV. See **Supplementary Video 2** for non-filtered data.

Previous mesoscale functional imaging studies on carnivore and primate V1s reported patchy spatial patterns of spontaneous activity that corresponded with the patchy patterns of iso-orientation columns (*3, 15, 22*). Using widefield calcium imaging, we first examined spontaneous activity across cortical areas and determined whether patchy spatial patterns of spontaneous activity exist beyond V1. The large sized FOV allowed us to reveal large-scale spatial patterns of spontaneous activity that extend across the cortical areas (*37, 38*). Importantly, the high spatiotemporal resolution of calcium imaging revealed patchy spatial patterns embedded within the large-scale patterns (**Supplementary Video 1, Supplementary Video 2**). Notably the patchy pattern was detectable throughout the FOV well beyond V1 including regions near parietal association areas (**Fig. 1d**), suggesting that the patchy spatial pattern is homogeneously present throughout the primate neocortex. Consistent with this idea, imaging in the temporal cortex revealed the presence of wave-like spontaneous activity with patchy texture (**Fig. 1e-g**; **Supplementary Video 3**), and even in the parietal and frontal areas where the presence of columnar functional map is not known (**Extended Data Fig. 1**). Collectively, these results together suggest that the patchy activity pattern is a canonical mode of spontaneous activity in the primate neocortex.

We performed principal component analysis (PCA) on the widefield spontaneous activity and then obtained a principal component (PC) that best correlated with the visually evoked activity to examine potential differences in spontaneous patches across areas, similar to the previous studies (e.g. (*39*)), using data covering V1 and the secondary visual cortex (V2) in the same FOV. In all three examined datasets, we found that the half-width-half-maximum of the autocorrelation profile was larger for V2 than for V1 (mean difference, 19.9%). Thus, the spontaneous patches in V2 were larger than those in V1 and similar in size to the orientation columns in each area (**Extended Data Fig. 2**), suggesting a potential correspondence between the spontaneous patches and functional columns. Cellular-scale imaging has also revealed differences in the spatial clustering of spontaneously coactive neurons across areas. We found that the proportion of coactive neurons in spatial proximity was significantly lower in the MT than in the V1 and V2 (*p* < 0.0001, two-sample *t* Test corrected by Bonferroni’s method; **Extended Data Fig. 3**), suggesting that spontaneously co-active neurons were less clustered in the MT than in the V1 and V2. These results suggest that spontaneous activity was heterogeneous across areas, based on the size of the meso-scale patchy organization and the degree of clustering of co-active neurons within the patches.

We next conducted two-photon imaging to compare spontaneous and stimulus-evoked activity at the cellular scale (**Fig. 2a**), and investigated whether the trial-averaged evoked responses were similar to the spontaneous patterns. Further, to address the question of across areal generality and potential species-related differences, we conducted the experiment on V1, V2 and the middle temporal visual area (MT) in the dorsal visual pathway of marmosets (*40*) and on mouse V1.

**Figure 2.**
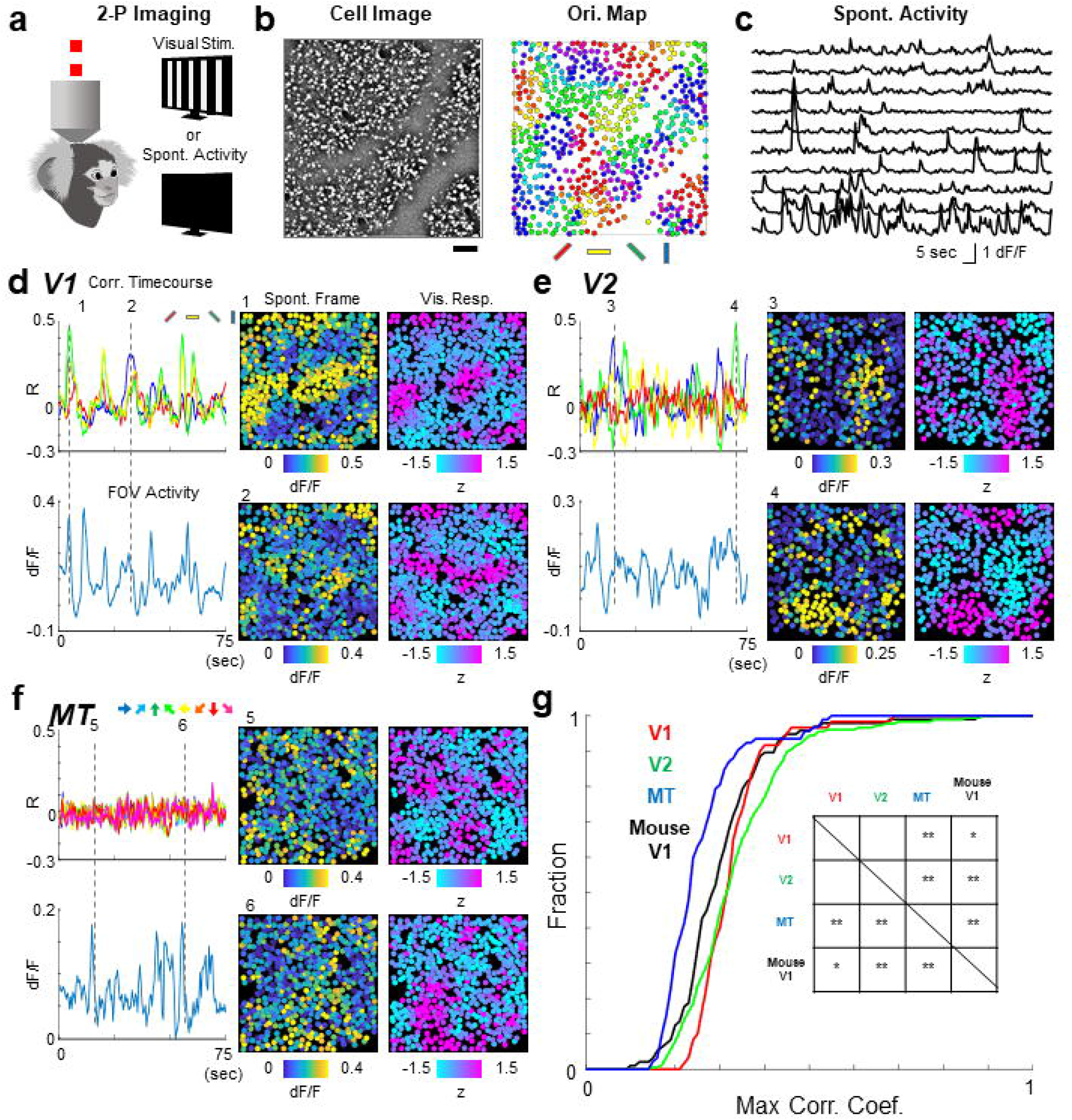
Spontaneous activity patterns were similar to evoked activity patterns in lower but not higher visual areas. **a)** Two photon calcium imaging were conducted in visual areas identified from widefield imaging. In each FOV, spontaneous and visually evoked activity were recorded for the identical set of cells. **b)** Example cell image and cellular orientation map in V1. Cellular orientation map was calculated using the cells identified from the cell image (left). Scale bar, 100 μm. **c)** Time courses of spontaneous activity for example cells. **d-f)** Comparison of spontaneous and evoked activity in V1 (d), V2 (e), and MT (f). Panel “Corr. Timecourses” shows correlation between individual spontaneous frames and responses to single orientations. Panel “FOV Activity” shows average activity of all the cells in the FOV. Panels “Spont. Frame” and “Vis. Resp.” show example spontaneous frame and a response to single orientation that was most similar to the spontaneous frame. Two examples are shown for time points indicated by dotted lines in the timecourses. **g)** Cumulative plots showing similarity of spontaneous frames and evoked activity. For each spontaneous frame, correlations with all single orientation responses were calculated, then the maximum value was taken. Inset, summary table of statistically significant difference [*, *p* < 0.05 (uncorrected), **, *p* < 0.05 (corrected by Bonferroni’s method), Wilcoxon’s Rank Sum test].

Consistent with previous reports on cat and macaque V1s (*3, 15, 39*), snapshots of spontaneous activity often appeared similar to visually evoked orientation responses (**Fig. 2b-d**). A similar correspondence between spontaneous activity and the functional map was also observed in V2 (**Fig. 2e**). However, in the downstream visual areas MT, we found much lower similarities of spontaneous activity and visually evoked orientation and direction responses, respectively (**Fig. 2f**; **Extended Data Fig. 4**).

Population data showed that, in all three areas, spatial correlation between single frames of spontaneous activity and evoked responses was high in V1 and V2, and became smaller in MT (**Fig. 2g**; **Extended Data Fig. 5a**). Widefield imaging also revealed that the similarity between spontaneous PCs and visually evoked maps was significantly higher in V1 and V2 than in the MT (*p* < 0.02, t-test; **Extended Data Fig. 6**). These results confirmed the previous finding of close correspondence between spontaneous and evoked activity in V1 (*3, 15-17, 22, 39*), and extended it to V2. However, the correspondence was not applicable to downstream area MT, suggesting that the relationship between spontaneous and evoked activity varies across cortical areas.

Subsequently, to obtain a better picture of the relationship between spontaneous and evoked activities across the three cortical areas, we next conducted geometrical analysis introduced in the recent mouse studies (*4*). We applied the geometrical analysis to both the marmoset data and also to the mouse data acquired in house using the same experimental techniques. Briefly, spontaneous and evoked neural population activities were expressed as vectors that were then projected onto three orthogonal subspaces within the activity space (**Fig. 3a**). The first subspace is shared between evoked and spontaneous activity (“shared space”). The second and third subspaces are respectively specific to spontaneous and evoked activity (“spontaneous-only space” and “stim-only space”). After projection to three orthogonal subspaces, the principal components (PCs) of each activity space were calculated to visualize the dominant activity patterns (**Fig. 3b; Extended Data Fig. 7**). As expected from the frame-by-frame correlation analysis, spatial similarity between PCs of shared space and visually evoked responses were high in V1 and V2 but low in MT (**Fig. 3c**, middle; see also **Extended Data Fig. 5b**). In contrast, the similarity between PCs of stim-only space and evoked responses were low in V1 and V2 but high in MT (**Fig. 3c**, top). PCs in spontaneous-only space were dissimilar to the evoked maps in all areas, as expected (**Fig. 3c**, bottom).

**Figure 3.**
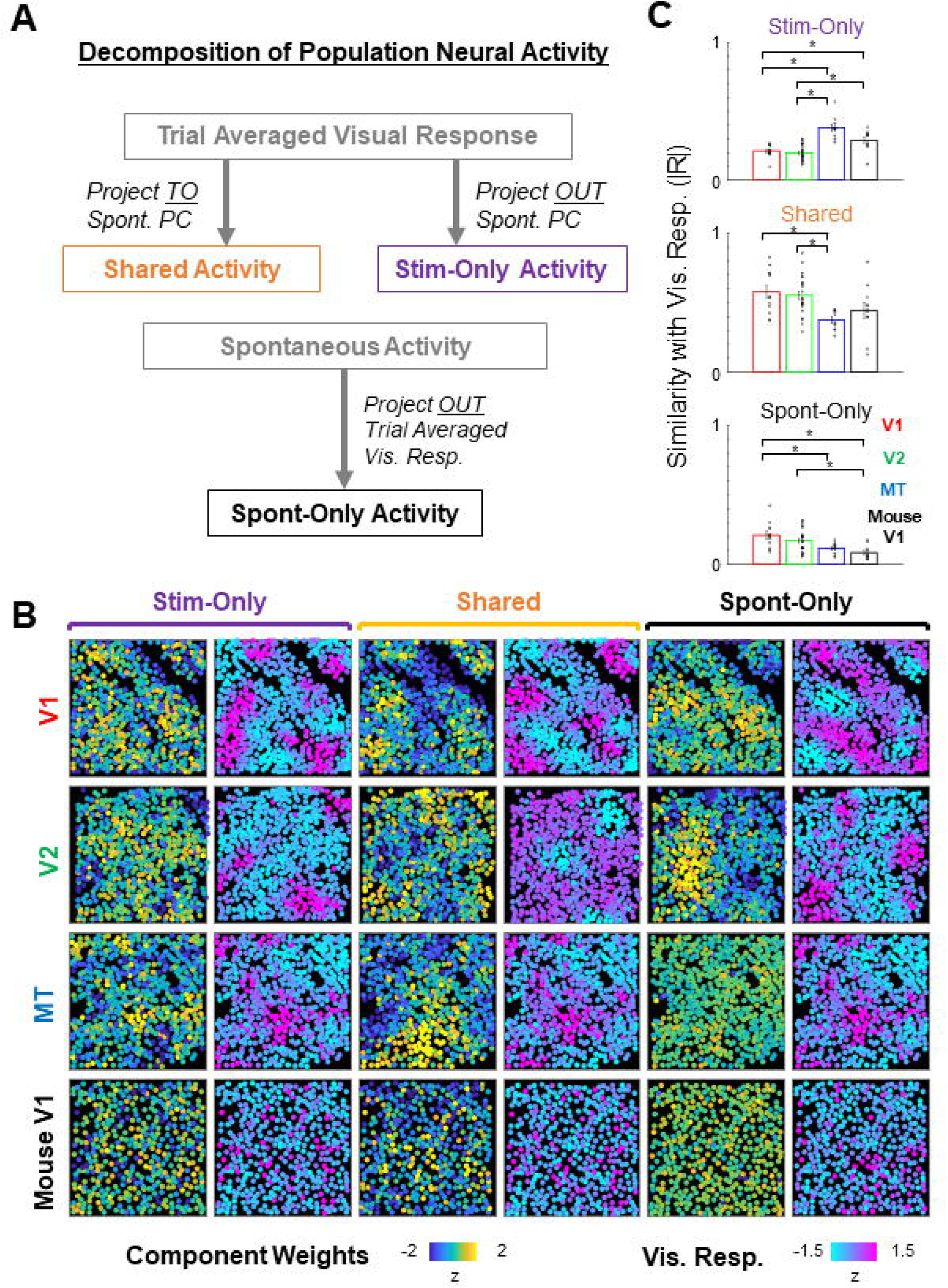
Decomposition of neural population activity to orthogonal subspaces. **a)** Overview of the analysis based on cellular-level neural activity data recorded by two-photon imaging. Trial averaged visual responses were projected to a space spanned by spontaneous activity to obtain activity patterns shared by spontaneous and evoked activity (Shared Activity). Trial averaged visual responses were orthogonalized to the space spanned by spontaneous activity to activity patterns specific to stimulus-evoked activity (Stim-Only Activity). Spontaneous activities were orthogonalized to a space spanned by trial averaged visual responses to obtain activity patterns specific to spontaneous activity (Sponta-Only Activity). **b)** Examples of 1^st^ principal components (PCs; left panels, blue-to-yellow color) of activity patterns in the three activity-spaces. For each PC, single condition visual response that was most similar is also shown (right, cyan-to-magenta color). Correlation coefficients between pairs of PC and visual response: V1 Shared (0.29), V1 Stim-Only (0.20), V1 Sponta-Only (0.13), V2 Shared (0.19), V2 Stim-Only (0.64), V2 Sponta-Only (0.13), MT Shared (0.12), MT Stim-Only (0.53), MT Sponta-Only (0.054), Mouse Shared (0.18), Mouse Stim-Only (0.64), Mouse Sponta-Only (0.057). **c)** Population data of the similarity between PCs and visual responses. Horizontal lines indicate pairs with significant difference (*p* < 0.05, two sample *t* tests, corrected by Bonferroni’s method for 6 pairwise comparisons). Error bars, SEM.

To quantify the contribution of the averaged signals to the total variance in neural activity, we performed a cross-validated regression of neural activity from the activity prediction. In the marmosets V1, V2, MT and the mouse V1, percent of the total variance explained by the averaged signals ranged between 30%–50% (**Extended Data Figure 8**). Thus, the trial-averaged neural activity explained a substantial fraction of the single-trial stimulus-evoked responses in all tested visual areas.

Finally, we focused on single-trial activities to quantify the fraction of variance of the visually evoked activity projected to the shared activity space (**Fig. 4a-b**). A previous study on mouse V1 reported that the fraction of the projected variance was approximately 20%, which led to the conclusion that stimulus-evoked activity and spontaneous activity are nearly orthogonal (*4*). In contrast to mouse V1, the fraction of stimulus-related variance explained by Shared Space was approximately 50% in the marmoset V1 (50 ± 2.7%, mean ± SEM). The projected variance slightly decreased in V2, but remained high (44 ± 2.3%). The projected variance decreased further in MT (31 ± 1.4%). The fraction of stimulus-related variance in the shared space for mouse V1 recorded using the same experimental setup yielded the smallest value (18 ± 2.0%), which is comparable to a previously reported value (*4*). Each PC in marmoset V1 and V2 had larger projected variances compared to marmoset MT and mouse V1. This was consistent with the relatively large, shared subspace in marmoset V1 and V2 compared to the other two areas (**Extended Data Fig. 9**). The same tendency was observed when shared space was limited to one dimension (**Extended Data Fig. 10**). The projected variance of the mouse V1 was similar after controlling the potential SNR differences between marmoset data and mouse data (**Extended Data Fig. 11**). The projected variance in marmoset V1 was significantly larger than that in mouse V1 after controlling for the number of cells and trials included in the analysis (**Extended Data Fig. 12b-c**). The same tendency across the three marmoset visual areas and mouse V1 was also observed when the data were summarized by animal (**Extended Data Fig. 5c**) and when non-trial averaged visual responses were used (**Fig. 4c**).

**Figure 4.**
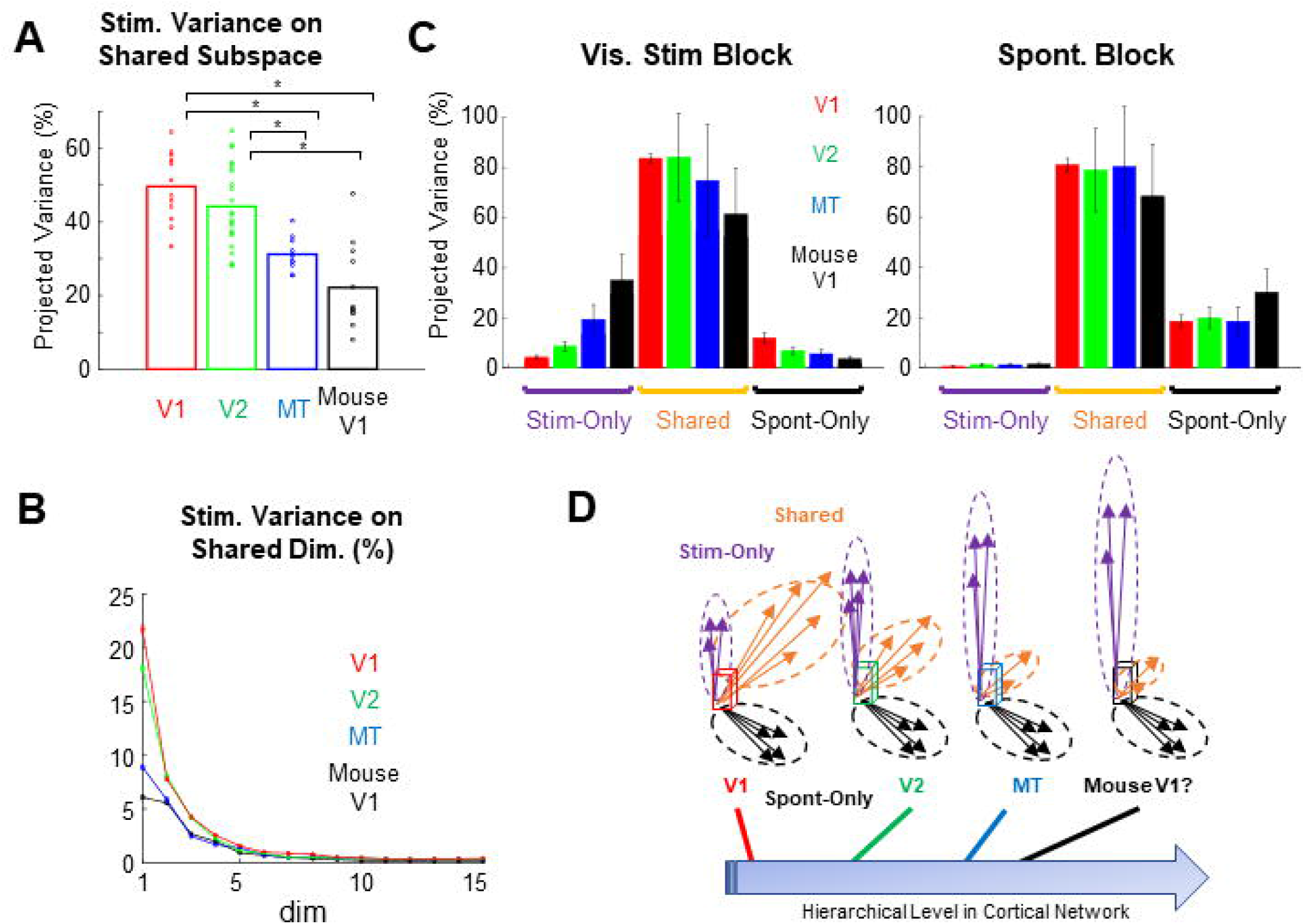
Progressive decrease of shared activity along the cortical hierarchy. **a)** Variance of cellular-level trial-averaged visual responses projected to the Shared subspace. Horizontal lines indicate pairs with significant difference (p < 0.01, two sample *t* test, corrected by Bonferroni’s method). Error bars, SEM. **b)** Same as (a) but projected to individual PCs of Shared Space. **c)** Variance of single trial visual responses (left) and spontaneous activity (right) projected to each subspace. Error bars, SEM. **d)** Schematics of the findings. Spontaneous and visually evoked activities in the primate visual cortex are progressively orthogonalized along the hierarchical network. Degree of the orthogonality in the mouse V1 is similar to higher but not lower visual areas of the marmoset.

The estimated dimensionality of shared space, as quantified by the number of PCs with large projections of stimulus-related variance, showed a similar tendency to the amount of projected variance; the dimensionality of shared space was larger for marmosets V1 and V2 than for marmoset MT and mouse V1 (**Extended Data Fig. 12a**). Collectively, these results suggest that, in the marmoset, the overlap between spontaneous and evoked activity progressively decreased along the cortical hierarchy.

## Discussion

In this study, we systematically investigated the patterns of spontaneous neocortical activity and its relation to stimulus-evoked activity across multiple hierarchically organized areas in the marmoset neocortex using mesoscale and cellular scale functional imaging with a newly developed expression system of genetically encoded calcium indicators in the primate brain. We found that spontaneous activity in the marmoset neocortex appeared as propagating waves with patchy spatial patterns on top of the waves. Notably, patchy spontaneous activity was not a unique feature of V1, but rather a general characteristic of primate neocortex, which includes the occipital, parietal, temporal, and frontal areas. However, the similarities between spontaneous and stimulus-evoked activity patterns varied across areas. In V1, the similarity between spontaneous and evoked activity was high, which is consistent with the findings of previous studies on cats and macaques. However, cellular imaging revealed lesser patchiness in the higher visual areas. Accordingly, stimulus-evoked and spontaneous activity were dissimilar in higher visual areas of the marmoset, as in mouse V1. The analysis of neural activity space further revealed that the stimulus-related variance projected onto the shared space progressively decreased along the cortical hierarchy (**Fig. 4d**). Thus, these results suggest that the marmoset visual network achieves the separation of stimulus-related signal information and internally generated noise through processing in the hierarchical cortical areal network.

Consistent with findings of previous studies on primates, carnivores, and rodents (*37, 38, 41*), we found that spontaneous activity appeared as propagating waves. Previous studies, particularly seminal works by Grinvald and colleagues, have also reported that spontaneous cortical activity in the primary visual areas of carnivores and primates show characteristic patchy spatial patterns (*3, 22, 39*). In the present study, we therefore examined the generalizability of the previous observations beyond the level of V1. The high signal-to-noise ratio of the primate-optimized calcium imaging allowed us to further reveal that patchy spatial patterns were superimposed on top of these waves. Notably, we revealed that patchy spontaneous activity was not a unique feature of V1, but rather a general characteristic of primate neocortex. In the cat and primate V1, iso-orientation columns are connected by horizontal connections (*42-44*). In the higher cortical areas of primates, both long-range and short-range cortico-cortical connections are known to have patchy spatial patterns in primates (*45, 46*). Thus, the presence of patchy spatial patterns may indicate that spontaneous activity propagates along cortico-cortical connections.

The present study confirms previously reported similarities between spontaneous and visually evoked activity in V1 of cats and macaques (*3, 15, 39*). Using the same recording method in mice, we also confirmed that state spaces spanned by stimulus-evoked responses and by internally generated noise were already orthogonal in V1 (*4, 5*). These findings showed that the apparent contradictory results in rodents and non-columnar mammals are not attributable to methodological differences (*47*) but rather to species-related difference in V1. The species-related difference may reflect the presence and absence of columnar cortical circuits in marmosets and mice, respectively (*35, 48*). Synaptic inputs to a single neuron are more likely to share similar function (*49*) in marmoset V1 with functional columns than in mouse V1 without functional columns (*50*). Because neurons with similar functional properties tend to have high noise correlation (*51*) as well as spontaneous correlation (*52*), marmoset V1 neurons are more likely to be activated by correlated inputs from presynaptic neurons that share similar functional properties than are mouse neurons. (**Supporting Discussion 1**). Recent studies on recurrent neural networks suggest that such differences in local connectivity patterns are related to the dimensionality of spontaneous and evoked neural network activity (*53, 54*). Avitan et al. reported that, in the zebrafish optic tectum, spontaneous and visually evoked activity patterns become less similar and geometrically diverse during development (*55*). It is of great interest for the future study to examine whether the relationship between the spontaneous and evoked activity in the mouse V1 and the marmoset MT follows a similar developmental pattern.

Of note, the use of anesthesia can affect spontaneous and stimulus-driven activity patterns. Okun et al. recorded spontaneous and evoked activities in the primary auditory cortex (A1) of anesthetized rats and, unlike the Stringer et al. study conducted in awake V1, showed considerable similarity between spontaneous and evoked neuronal activity patterns (*56*). Thus, the use of anesthesia can affect the degree of similarity between spontaneous and evoked neuronal activities. In the present study, to account for this possibility and minimize the effect of anesthesia on the comparison between mouse V1 and marmoset V1, we conducted mouse experiments under anesthetized conditions comparable to those used for marmosets. Hence, the difference between marmoset V1 and mouse V1 found in the present study is unlikely to be explained solely by anesthesia. However, some differences in the study by Stringer et al. and the present study in mouse V1 (e.g. larger dimensionality of the shared space) may be attributable to the use of anesthesia. Stringer et al. reported that the one-dimensional behavior-stimulus shared subspace can be interpreted as a multiplicative gain modulation of cortical stimulus responses (see their Fig. 4L). Such multiplicative gain modulation has been linked to attentional modulation of cortical activity (*57, 58*). It is likely that the anesthetic condition used in the present study strongly attenuated this component of neuronal activity (in fact the variance projected to the 1st PC in our data was 8% whereas that in Stringer et al. was 15%).

Previous studies on information processing in the hierarchical cortical network reported increasingly larger spatial scales (*i*.*e*., receptive fields) (*59*) and temporal scales along the cortical hierarchy (*60*). In this study, we revealed a novel computational role for the hierarchical cortical network in progressively orthogonalizing spontaneous and stimulus-evoked activity. Such orthogonalization likely continues to the frontal cortex, where neurons form distinct assemblies in spontaneous and task-related periods (*61*). A potential functional role of orthogonalization may be the separation of stimulus-related signals and internal noise in the neural activity space (*5, 7, 8, 11, 12, 62-64*). Importantly, our study showed that, unlike orthogonalization in mice, which is already completed in V1, that in marmosets takes place at the multiple steps of interareal processing. One possible explanation for the progressive orthogonalization is that the activity patterns of stim-only PCs are more effective in activating neurons in the downstream area than those of shared PCs. Indeed, Semedo et al. reported that V2 activity reflects specific patterns of V1 activity but not necessarily the largest activity (*65*). Such across areal activity transmission filters out shared PCs and hence, enables segregation of sensory signals and internal noise (**Supporting Discussion 2**). Because almost all of the visual areas are accessible for optical recording and manipulation, the marmoset monkey is ideally suited for testing these hypotheses.

The similarity of spontaneous and evoked neuronal activity has been interpreted as evidence supporting the functional significance of spontaneous cortical activity, such as the Bayesian prior (*17*), recent history of experience (*66*) and prediction of incoming inputs(*67*). In marmoset V1, the estimated number of dimensions in the shared subspace was three, which was larger than the dimensionality of mouse V1, but not by a large margin (**Extended Data Fig. 12a**). These results suggest that even if spontaneous activity in marmoset V1 serves as Bayesian prior, it may only provide rudimentary information about the visual scene (e.g. orientation or color). Nevertheless, the overlap of spontaneous and evoked activity patterns found in marmoset V1 is consistent with the hypotheses about the functions of spontaneous activity listed above. In contrast, in mouse V1, spontaneous and sensory-evoked activities are nearly orthogonal and represent behavioral information such as facial expression (*4*). The present study shows that the marmoset visual areas downstream to V1 achieve an orthogonal relationship of spontaneous and sensory-evoked activities, suggesting that spontaneous activity in these areas may represent behavioral information (*68*), similar to mouse V1. Interestingly, a recent study suggested that neural coding in V1 and V2 was optimized for sensory representation, whereas that in the MT was optimized for visual discrimination (*69*). Ruff and Cohen have reported that visual attention further optimizes stimulus representation by MT neurons by aligning it to the axis of the activity space, which is important for guiding behavior (*70*). It is tempting to speculate that the difference of the site of orthogonalization in the marmoset and the mouse is related to the difference of the depth of the hierarchical cortical network in the two species: the marmoset with a deep cortical network may benefit initially parallel spontaneous activity, whereas the mouse with a shallow cortical network benefit more by having orthogonalized spontaneous activity already at V1. Nevertheless, despite the species difference in circuit-level implementations, the orthogonalization of activity patterns is likely a common computational goal for separating sensory signals and internal noise in biological neural circuits.

## Methods

### Animals

A total of 12 adult common marmosets (*Callithrix jacchus*; 7 males and 5 females; body weight, 280-350 g; age, 1-2 years) obtained from Nihon Clea and three Thy1-GCaMP6 mice (4 males; GP 4.3, JAX#024275; body weight, 28-35 g; age, P50-60) obtained from Jackson Laboratory were used. All animals were housed in a 12:12 h light-dark cycle, had access to water and food *ad libitum* and were not used for other experiments prior to the present study. All animal experiments were carried out in accordance with the institutional welfare guidelines laid down by the Animal Care and Use Committee of the University of Tokyo, and approved by the Ethical Committee of the University of Tokyo. Of the 12 marmosets, 10 were used to obtain widefield imaging data [Large FOV covering the occipital-parietal cortex, two animals; V1/V2, three animals; MT, four animals; Frontal cortex, one animal], and nine were used to obtain two-photon imaging data [V1, 12 FOVs in three animals; V2, 23 FOVs in four animals; MT, 11 FOVs in four animals]. Four mice were used to obtain the two-photon imaging data (12 FOVs in V1).

### Plasmid Construction

pAAV-Thy1S-tTA and pAAV-TRE-GCaMP6f-WPRE were kindly provided by Dr. T. Yamamori (*18*). All newly designed GCaMP expression plasmids were constructed using pAAV-TRE-GCaMP6f-WPRE as a template. To generate pAAV-TRE-GCaMP6f/s-P2A-GCaMP6f/s-WPRE, GCaMP6s and P2A gene fragments were obtained from pGP-CMV-GCaMP6s-WPRE (Addgene: #40753) and pAAV-hSyn1-GCaMP6s-P2A-nls-dTomato (Addgene: #51084), respectively.

### Virus Production

Details of the production of primate-optimized GCaMP and adeno associated virus (AAV) are described in detail elsewhere (Hashimoto et al., in preparation). Briefly, AAV plasmids were packaged into AAV serotype 9 using the AAV Helper-Free system (Agilent Technologies). pAAV vector, pRC9 and pHelper plasmids were transfected into HEK293 cells. At 72 hours after transfection, AAV2/9 particles were purified using the AAV Purification kit (Takara). The AAV solution was concentrated to the optimal volume (up to 100 μL) by centrifugation using an Amicon Ultra-4 100k centrifugal filter unit (Millipore). To quantify the virus titer, the number of genomic copies was quantified with intercalating dyes (Thermo Fisher Scientific) and two sets of primers for WPRE or hGHpA genes using a LightCycler 480 (Roche). The final titration of AAV was estimated as relative quantitation based on the calibration curve calculated from the known number of AAV plasmids copies.

### Virus Injection

All surgical procedures were performed under aseptic conditions. Marmosets were anaesthetized with isoflurane (4.0-5.0% for induction and 1.5-3.0% for maintenance in a mixture of 20-50% O_2_ and air). Throughout surgery, percutaneous oxygen saturation (SpO_2_) and heart rate were monitored and maintained at > 96% and < 200 bpm, respectively. The rectal temperature was maintained at 37 °C using an electric blanket and a feedback-controlled heating pad. Before surgery, cefovecin (14 mg/kg, i.m.) or ampicillin (15-20 mg/kg, s.c.) was administered as an antibiotic prophylaxis. Meloxicam (0.1-0.2 mg/kg, s.c.) was administered to reduce pain and inflammation. The scalp was sterilized with povidone iodine. All incision sites were pre-treated with local injections of lidocaine HCl (2%) or lidocaine jelly. After scalp incision, a custom-made metal head post was attached to the skull using dental acrylic (Shofu Inc.). Marmosets were then head-fixed on a custom-made metal stage using a head post.

For AAV injections, small craniotomies (1-2 mm in diameter, 4-12 sites, 1-2 mm apart) were made around the targeted cortical regions, and the dura mater was cut to expose the cortical surface. The brain was washed with gentamycin-mixed artificial cerebrospinal fluid (ACSF). An AAV cocktail with a 1:1 mixture of AAV2/9-Thy1S-tTA (3.11×10^13^ vg/ml) and AAV2/9-TRE-tandem-GCaMP6s (2.65×10^13^ vg/ml) was loaded into a glass pipette and injected using a Nanoject III (Drummond Scientific Company). The pipette was inserted 300-500 μm below the cortical surface, and 0.5-1.0 μl AAV was injected at 0.12 μl/min in a single injection site. After the injection, the craniotomies were covered with Kwik-Sil™ (World Precision Instruments), the head post was detached, and the scalp was sutured. All wound sites were treated with gentamycin ointment. After the surgery, warmed physiological saline (5 ml) was administered subcutaneously to prevent dehydration, and the animals were returned to their home cage for recovery. The next day, meloxicam and/or ampicillin were/was administered for postoperative management. The animals were maintained for 4-5 weeks before the imaging experiments to ensure sufficient GCaMP expression.

### Preparation for Imaging Experiments

The marmosets were treated with atropine sulfate (0.2 mg/kg, i.m.) and anesthetized with a mixture of ketamine (45 mg/kg, i.m.) and xylazine (3.0 mg/kg, i.m.). SpO_2_, heart rate, and rectal temperature were monitored as previously described (*71*) (Hashimoto et al., in preparation). Meloxicam (0.1-0.2 mg/kg, s.c.) was administered as an anti-inflammatory drug. A tracheotomy was performed, and the animal was mechanically ventilated. End-tidal CO_2_ was monitored and kept at 3.4-4.0% throughout the experiment. An intravenous catheter was placed in the femoral vein, and anesthesia was maintained by constant infusion of remifentanil (6.0-15.0 μg/kg/h, i.v.) mixed in lactated Ringer’s solution (2.0 ml/kg/h) and dexamethasone (0.4 mg/kg/h). Muscle relaxation was induced by vecuronium bromide (0.1 mg/kg/h, i.v.). Additional doses of isoflurane (2.0-3.0%) and N_2_O (50% in O_2_ and air) were administered intraoperatively. A custom-made metal head post was attached to the skull, and the animal was mounted on a stereotaxic apparatus. A large cranial window was made over the targeted brain regions (12-mm diameter circle or 8 mm × 16 mm square, depending on the targeted brain regions), and the dura mater was opened. A glass coverslip (10 mm in diameter or 8 mm × 16 mm square) attached to a custom-made metal rim was placed on the exposed cortical surface, and the brain was sealed with Kwik-Sil™ (World Precision Instruments) and dental cement (Sun Medical). During imaging, anesthesia was maintained by remifentanil (6.0-15.0 μg/kg/h, i.v.), and doses of isoflurane and N_2_O were decreased to 0-0.5% and 10-50%, respectively. The eyes were covered with contact lenses and frequently moistened with an ophthalmic solution to ensure a clear view throughout the experiment.

### Widefield Ca^2+^ Imaging

*In vivo* wide-field imaging was performed using a macrozoom fluorescence microscope (MVX10, Olympus) equipped with a 2× objective (2x MVX Plan Apochromat Lens, NA 0.25, Olympus). GCaMP was excited by a mercury lamp through a GFP mirror unit (U-MGFPHQ/XL, Olympus, excitation peak: 488 nm, emission peak: 507 nm). Fluorescence images were acquired using an sCMOS camera (Zyla 4.2 sCMOS, Andor Technology) controlled by NIS Elements BR (Nikon). A square region of the cortex (6 ×□ 6 mm to 15 ×□15 mm, 512□×□512 or 256□×□256 pixels) was imaged at 5 Hz.

### Two-Photon Ca^2+^ Imaging

After widefield imaging, the animals were moved under a two-photon microscope (A1RMP, Nikon, Tokyo, Japan) equipped with a water immersion objective (16× or 25× with NA 0.8 or NA 1.1, respectively, Nikon) to perform two-photon calcium imaging. GCaMP was excited at a 920-nm wavelength using a Ti:sapphire laser (Mai Tai HP DeepSee, Spectra Physics). A square region of the cortex (800 μm × 800 μm in 512 × 512 pixels with a 16× objective for marmosets; 800 μm × 800 μm in 512 × 512 pixels with a 25× objective for mice) was imaged at 2 Hz. The position of the FOV was selected from the functional map obtained using widefield imaging. The spatial pattern of blood vessels on the cortical surface was used to guide the FOV. The depth of the imaged plane was carefully adjusted manually between the scans. Image planes from the same cortical location were separated by at least 30 μm in depth to avoid imaging the same neurons twice.

### Visual Stimulation

Visual stimuli were generated using custom-written programs in PsychoPy (*72*). A 32-inch LCD monitor with a 60-Hz refresh rate (ME32B, Samsung) was positioned 30 cm in front of the marmosets. The stimulus screen spanned –35.6° to 35.6° horizontal and –50.7° to 50.7° vertical of the animal’s visual field. The stimuli were presented to both eyes. For mapping orientation preferences, a drifting square-wave grating [100% contrast; 0.5 to 1 cycles per degree (cpd); 2 Hz] tilted at one of four to eight orientations in equal steps, moving in one of two directions orthogonal to the orientation was presented. To map orientation preferences, we presented a drifting square-wave grating [100% contrast; 0.5–1 cpd; 2 Hz] tilting at one of four to eight orientations in equal steps and moving in one of two directions orthogonal to the orientation. For mapping direction preferences, random dots (100% coherence; dot size, 0.5 °; speed, 20 °/s] moving in one of eight or twelve directions in equal steps were used. Each stimulus started with a blank period of uniform gray (4s) followed by the same period of visual stimulation. Each condition was repeated 20–100 times in pseudorandom orders.

In both the widefield and two-photon imaging experiments, spontaneous activity was recorded separately from the visually evoked responses in the dedicated scans. During scans for spontaneous activity recordings, the LCD monitor used for visual stimulation was turned off. For each FOV, we conducted a spontaneous activity scan either before or after the visual stimulus scans. The interval between the two scans was no longer than 30 min.

### Widefield Imaging Data Analysis

All analyses were performed using custom-written programs in MATLAB (MathWorks, Natick, MA, USA). For widefield imaging data, small drifts between the imaging frames were realigned by maximizing the correlation between the images. The relative change in fluorescence (Δ*F*/*F*) was computed using the following equation: Δ*F*/*F* = (*F* – *F*_0_)/*F*_0_. For data with visual stimulation, *F*_0_ is the average fluorescence during the pre-stimulus period (baseline), and *F* is the fluorescence during stimulus presentation. For spontaneous activity data, *F*_0_ is the average across the entire scan, and *F* is the fluorescence during spontaneous activity. To compute trial-averaged responses, images or time courses were sorted by stimulus conditions and averaged across all repetitions.

The method for calculating orientation preference maps and direction preference maps has been described elsewhere (*71*). Briefly, single-condition maps to the tested orientation (or directions of motion) stimuli were obtained by calculating the relative change in fluorescence (Δ*F*/*F*) between the baseline (1 s before stimulus onset) and the stimulus period (4 s). Each map was high-pass filtered by subtracting the low spatial frequency background, which was obtained by applying 2D median filters three times (kernel size: 0.45 mm by side) to the original map. Direction preference maps (HLS maps) were obtained from single-condition maps. In the HLS map, hue (H) represents the preferred orientation or direction calculated by vector averaging, saturation (S) represents the global measure of orientation or direction tuning, which corresponds to 1 – CV (circular variance in orientation or direction preference), and lightness (L) represents the response to the best direction (Δ*F*/*F*).

Movies of spontaneous activity are shown in Δ*F*/*F*, calculated as described above. For our analysis of patchy spontaneous activity in widefield imaging, we followed procedures commonly employed in previous studies, namely spatial filtering and PCA (*22, 39*). We first down-sampled the images from 512 × 512 pixels to 256 × 256 pixels, followed by spatial highpass filtering (σ = 180 μm for occipital-parietal FOV, and σ = 280 μm for V1/V2 and MT FOV). We consider that the use of high-pass filtering did not artificially create patchy spatial patterns; first, we used high-pass filters instead of bandpass filters. Whereas bandpass filters may artificially bias patchy patterns by selectively suppressing both low and high spatial frequency components, high-pass filters are less likely to have such a bias toward a particular frequency band. Second, patchy spatial patterns that resemble the spatial patterns of the orientation columns are readily visible in the data without high-pass filtering (Supplementary Videos 1-3). To obtain underlying spatial patterns from the data in an unbiased manner, PCA was then applied to the filtered images. PCs clearly showing the spatial patterns of blood vessels were discarded (typically three to five PCs in the first 20 PCs; see **Extended Data Fig. 13** for the top 20 PCs for example data). To estimate the spontaneous patch size (**Extended Data Fig. 2**) for each FOV, we first searched for a spontaneous PC that best correlated with the visually evoked activity. We chose to analyze widefield imaging data covering V1 and V2 in the same FOV because these data allowed us to compare the two cortical areas without being affected by differences in animals and/or other experimental conditions. Autocorrelation analysis was performed separately for V1 and V2, and the sizes of the two autocorrelation profiles were compared. Because this analysis used widefield imaging data obtained from one FOV per animal, we did not have sufficient data to perform statistical testing.

### Two-photon Imaging Data Analysis

For the cell-level analysis, cell bodies were automatically identified by template matching with a circular template using high-pass filtered images of the temporal variance of fluorescence signals (Fig. 2b). Time courses of individual cells were extracted by summing pixel values within the contours of cell bodies and then converted to Δ*F*/*F* using a percentile filter with a sliding window (baseline, lower 30 percentile; window size, 24.5 s), removing the slow drift of signals. For the data obtained from transgenic mice, we additionally subtracted background signals to minimize contamination of neuropil signals (*73*). For each cell, the background signal time course was obtained by averaging the pixels surrounding the cell body, and then, the background signal time course was subtracted from the cell’s time course after multiplying by a scaling factor (0.7). In a control analysis that accounted for a potential SNR difference between mice and marmosets due to the difference in magnification (**Extended Data Fig. 11**), we set the number of pixels allocated to each cell to 100 for both mice and marmosets.

Visual responses were calculated using the time-course averaged across trials (repetitions). The response to each orientation condition was defined as the mean of the responses to two drifting gratings moving in opposing directions orthogonal to the orientation. The response to each direction condition was defined as the mean of the responses to gratings or dots moving in the direction specified by the condition. The cellular orientation map (Fig. 2b) was obtained by calculating the preferred orientation for each neuron using vector averaging of the orientation responses (*71*). To compare visually evoked and spontaneous activity patterns, we selected FOVs (1) with stable recordings of both visual stimulation scan and spontaneous activity scan and (2) with sufficiently large number of active cells (> 100).

To analyze the clustering of spontaneously active neurons (Extended Data Fig. 3), we adopted the clustering index used in previous studies (*71, 74*). For each spontaneous frame, we calculated the median of absolute pairwise difference of spontaneous activity between pairs of neurons across all the pairs that were located within 400 μm (orientation column spacing in the marmoset V1(*35*)). Then, we did the same calculation for the position-shuffled control. The clustering index for the plane was defined as the value obtained for the position-shuffled control divided by the value obtained for the real data.

Neuronal activity was decomposed into orthogonal subspaces as described by Stringer et als (*4*). We specifically used the procedure described by Stringer et al. to analyze spontaneous activity, which did not use any behavioral information. Visual stimulation and spontaneous activity scans were obtained separately for each group of neurons within the same FOV. We projected the visually evoked and spontaneous activities into three orthogonal subspaces: stimulus-only space, shared space, and spontaneous-only space, using the following procedures (**Fig. 3a**). The fraction of single-trial stimulus-evoked response explained by the trial-averaged neural activity was estimated using cross validation. Half of the trials were used to make the predictor (i.e., trial-averaged neural activity), and the other half were used to estimate the trial-to-trial variance. In the analysis in which we controlled for the number of cells and trials (**Extended Data Fig. 12b-c**), we set the number of cells and trials to be equal across all FOVs by randomly selecting 400 cells and 20 trials in each FOV.

For each visual stimulation scan, neural activity was averaged across trials. We treated each frame of the trial-averaged visual evoked response as an individual visual response pattern (frames corresponding to the stimulus-off periods were discarded). Depending on the number of orientations/directions tested, the total number of trial-averaged visual responses for each set of stimuli ranged from 32 to 64. The frames in which visual stimuli were presented were then collected and denoted as trial-averaged visual responses, producing a matrix *M*_*vis*_ of trial-averaged visually evoked activity with dimension *N*_*neuron*_ × *N*_*stim*_. For cross-validation, we used half of the trials to obtain *M*_*vis*_, and spared the other half of the trials for later calculation of the projected variance of the visually evoked activity. Spontaneous activity *M*_*spont*_, which is a matrix *N*_*neuron*_ × *N*_*spont-frame*_, was subjected to PCA, and the first 50 PCs were retained 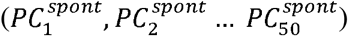. Note that, for each FOV, *N*_*neuron*_ was the same for the visually evoked activity and spontaneous activity. The stimulus-only activity *M*_*vis-only*_ was obtained by regressing out 50 spontaneous PCs from each column of the trial-averaged visual response, *M*_*vis*_. Thus, each column *e*_*i*_ in *M*_*vis-only*_ is orthogonal to the spontaneous PCs.

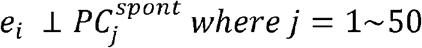

We set the number of spontaneous PCs to 50 for the following reasons. In the spontaneous activity scans, the number of frames (>1000) exceeded the number of cells (<1000). Therefore, the full dimensions of the spontaneous space are equal to the number of cells. Because the visual responses were recorded in the same set of cells, if we used whole spontaneous space, the entire variance of the visual responses would be projected onto the spontaneous space. Thus, we limited the spontaneous space to smaller dimensions. To set the number of PCs to use, we referred to a previous study by Stringer et al., where they used the first 128 dimensions of spontaneous PCs for the recording of ∼10000 cells. Because the number of cells recorded in our study was approximately 1000 at most (∼500 on average), we reasoned that 50 PCs would be sufficient to construct a shared space. To confirm that the present results did not depend on the exact number of PCs used, we conducted the same analyses with 100 spontaneous PCs (**Extended Data Fig. 14**). The matrix of shared activities *M*_*shared*_ was obtained by projecting each column *v*_*i*_ of the trial-averaged visual responses *M*_*vis*_ onto 50 spontaneous PCs. Thus, each column *w*_*i*_ of the shared activity *M*_*shared*_ is contained in the space spanned by 50 spontaneous PCs:

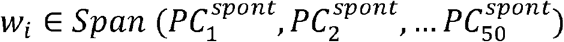

The matrix of spontaneous-only activity *M*_*spont-only*_ was obtained by regressing out each column *v*_*i*_ of the trial-averaged visual responses *M*_*vis*_ from each column (*i*.*e*., frame) *s*_*i*_ of spontaneous activity *M*_*spont*_. Thus, each column *u*_*i*_ of the spontaneous-only activity *M*_*spont-only*_ was orthogonal to each column *v*_*j*_ of the visually evoked activity *M*_*vis*_:

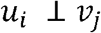

Because the number of trial-averaged visual responses was smaller than the number of recorded cells in each FOV (>100), visual response space did not encompass full neural activity space which has number of dimensions equal to the number of recorded neurons. Thus, we projected out all the trial-averaged visual responses instead of projecting out some PCs of the trial-averaged responses. The PCs of each subspace (**Fig. 3b**) were obtained by applying singular value decomposition or PCA, and the PCs of the shared space were obtained by finding a sequence of orthogonal directions *a*_*i*_ that maximized the sum of squares of the trial averaged visual response *M*_*vis*_:

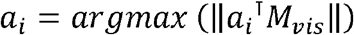

such that

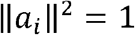

and

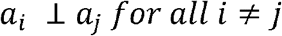

and

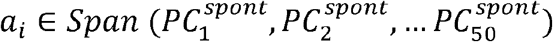

This was calculated as the left singular vector of the shared activity *M*_*shared*_. PCs of stim-only space were similarly obtained as the left singular vector of the stimulus-only activity *M*_*vis-only*_. The PCs of the spont-only space were obtained as the principal components of the spont-only activity *M*_*spont-only*_. PCs that explained less than 3% of variance were discarded.

The similarity between each PC and the visual responses (i.e. single-condition responses) was calculated using a cell-wise correlation. The amount of stimulus-related variance projected onto the shared space was calculated as the stimulus-related variance projected to the space spanned by the first 50 spontaneous PCs that were used to calculate the shared space. Trial-averaged visual responses *M*_*vis*_ were projected onto the first 50 spontaneous PCs 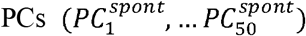, and the sum of the squared projection lengths was calculated (**Fig. 4a**). To quantify the amount of variance in the stimulus-related activity projected onto each shared PC, we used the held-out half of the visual stimulation trials, which were not used to calculate the shared PC. The trial-averaged responses calculated using the held-out trials were projected onto each shared PC, and the squared projection length was used as the projected variance on the shard PC (**Fig. 4b**).

For the projection of single trial activity to subspaces (**Fig. 4c**), we used PCs of the three subspaces obtained as described above (note that PCs with greater than 3 % explained variance were used). Neural activity in each frame of the held-out trials of the visual stimulation scans (with the blank period discarded) and spontaneous activity scans were projected onto the PCs of the three subspaces. For each subspace, the sum of the squared projected lengths was calculated and regarded as the amount of variance projected onto the subspace. The variances projected onto the three subspaces were calculated for each scan and normalized to the total variance, which was the sum across frames. Note that the projected variance on sponta-only PC and stim-only PC was non-zero in the visual stimulation block and the spontaneous block, respectively (Fig. 4C). Because single-trial visually evoked response not only contains visually evoked components but also non-visual (“spontaneous”) components in additive manner (*4, 6*), trial-to-trial variability of single-trial visually evoked responses in our data likely contained both visually related variability and spontaneous activity-related components. We believe that some of the latter components were projected onto the sponta-only PC. Regarding the non-zero projected variance on stim-only PCs in the spontaneous block, we believe that this was because we projected out top 50 PCs of spontaneous activity, instead of the full PCs, to define stim-only space. The residual PCs of spontaneous activity that were not used to define stim-only space could have contributed to the projection to the stim-only space.

### Statistics

Statistical tests were conducted using Statistics Toolbox in Matlab. Independent group comparisons were performed using two-sample Kolmogorov-Smirnov tests, Wilcoxon rank-sum test, and two-sample *t-tests*. No statistical methods were used to pre-determine sample sizes, but our sample sizes were similar to those used in previous studies. Allocation in the experimental groups was not randomized. Data collection and analysis were not blinded to experimental conditions.

## Supporting information

Supporting Discussion

## Acknowledgements

We thank members of the Ohki laboratory for helpful discussions; A. Hayashi, Y. Kato, M. Taki, T. Inoue, Y. Sono, T. Honda for animal care; Drs. T. Yamamori, A. Watakabe for providing pAAV-Thy1S-tTA and pAAV-TRE-GCaMP6f; Drs. T. Ebina, Y. Masamizu, M. Matsuizaki for technical advice; The Genetically-Encoded Neuronal Indicator and Effector (GENIE) Project for providing GCaMP6f/s.

## Author contributions

T. Matsui., T.H., T. Murakami and K.O. conceived the project. M.U. developed and provided GCaMP expression systems optimized for primates. T. Matsui. T.H., T. Murakami, M.U., K.K., and T.K. performed *in vivo* experiments. T. Matsui, T.H., T. Murakami, and K.O. analyzed the data. All authors discussed the results. T. Matsui, and K.O. wrote the manuscript with contributions from all authors. T.Matsui, T.H., T. Murakami, and M.U. contributed equally to this work.

## Reporting summary

Further information on research design is available in the Nature Research Reporting Summary linked to this paper.

## Data and materials availability

The datasets and codes used in this study are available from the corresponding authors on reasonable request.

## Extended Data Figure Legends

**Extended Data Figure 1** | **Examples of patchy spontaneous activity in the parietal and frontal areas. a-d)** Spontaneous activity in the FOV including the parietal areas. (a) The approximate position of the FOV on the marmoset brain atlas (*75*). (b) FOV image showing in vivo GCaMP fluorescence. (c)-(d) Snapshots of spontaneous activities. **e-h)** Snapshots of spontaneous activity in FOV including the frontal areas. (b) Approximate position of FOV. Same convention as Fig. 1a (b) FOV image showing in vivo GCaMP fluorescence. (c)-(d) Snapshots of spontaneous activity. Scale bars, 1 mm.

**Extended Data Figure 2** | **Size of spontaneous patches varied across cortical areas. a)** Left column, principal components (PCs) of spontaneous activity for a FOV covering V1 and V2. PC#3-7 are shown. PC#1 and PC#2 are discarded because these PCs mainly reflected vascular noise (see also Extended Data Fig. 13). Middle and right columns, spatial autocorrelation maps of PCs obtained using pixels in V1 (middle) and V2 (right). For all PCs, autocorrelation peaks in V2 were broader than those in V1. **b)** A visually evoked orientation response in the same FOV as (a). **c-d)** Quantification of the sizes of the autocorrelation peaks in V1 and V2. Data from three FOVs covering V1 and V2 were used (including a FOV shown in (a)). 5 PCs were obtained from each FOV. For each PC, autocorrelation peaks in V1 and V2 were fitted with two dimensional Gaussians. Lines in (c) and (d) connects datapoints taken from the same PC in the same FOV. Autocorrelation peaks were larger in V2 than in V1 for both longer (c) and shorter axes (d) (mean difference, 19.9%).

**Extended Data Figure 3** | **Clustering of cellular spontaneous activity differs across areas**. For each spontaneous frame, the degree of clustering of spatial clustering of active neurons was assessed (see Methods). Larger clustering index indicates more spatial clustering of active neurons. Degree of clustering decreased in higher visual areas (*, p < 0.001, Kolmogorov-Smirnov test, corrected by Bonferroni’s method; **, P = 0.0481, Kolmogorov-Smirnov test, uncorrected). Error bar, SEM.

**Extended Data Figure 4** | **Example cellular spontaneous and evoked activity in mouse V1**. Same convention as in Fig. 2d-f but for an example FOV in mouse V1 (b). Scale bars, 100 μm.

**Extended Data Figure 5** | **Results summarized by animals. a**) Same as Fig. 2g but data were summarized by animals. **b**) Same as Fig. 3c but data were summarized by animals. In **c**) Same as Fig. 4a but data were summarized by animals.

**Extended Data Figure 6** | **Comparisons of spontaneous PCs and visually evoked maps. a**) An example for a FOV in V1/V2. Left, an example of spontaneous PC that resembled orientation evoked response. Right, a map of preferred orientations. White and black contours indicate locations of the positive (yellow) peaks and negative (blue) peaks in the spontaneous PC, respectively. **b**) Same convention as (a) but for an example FOV in MT. Left, spontaneous PC. Right, a map of preferred directions. **b**) Spatial correlation between spontaneous PCs and orientation/direction maps. For each FOV, the highest correlation between orientation (V1/V2) or direction (MT) evoked response was calculated. The spatial correlation between spontaneous PCs and functional maps was significantly higher in V1/V2 than in MT (*p* < 0.017, two-sample t test). Scale bars, 1 mm.

**Extended Data Figure 7** | **Data related to PCA and SVD of cellular scale neural activity. a**) Plots of cumulative explained variance for PCAs which were used to obtain spontaneous PCs (top) and spontaneous-only PCs (bottom). Each trace indicates the cumulative explained variance for one FOV. **b**) Plots of singular values for SVDs which were used to obtain shared PCs (top) and evoked-only PCs (bottom). Each trace indicates the plot of singular values for one FOV.

**Extended Data Figure 8** | **Fractions of the variance of single-trial stimulus evoked activity explained by trial-averaged stimulus evoked activity**.

**Extended Data Figure 9** | **Stimulus-related variance projected onto each shared PC**.

**Extended Data Figure 10** | **Stimulus-related variance projected onto one-dimensional shared space**. Same convention as Figure 4a but projected variances were calculated using only the 1^st^ shared PCs.

**Extended Data Figure 11** | **Stimulus-related variance projected onto shared space with SNR adjusted estimate of the mouse V1**. For each spontaneous frame, the degree of spatial clustering of the active neurons was assessed (see Methods). A larger clustering index indicates greater spatial clustering of active neurons. The degree of clustering decreased in the higher visual areas (*, p < 0.001, Kolmogorov-Smirnov test, corrected by Bonferroni’s method; **, P = 0.0481, Kolmogorov-Smirnov test, uncorrected). Error bar, SEM.

Variance in cellular-level trial-averaged visual responses projected onto the shared subspace. V1, V2, MT, and Mouse V1 show the same data as in Fig. 4a. “Mouse V1 (SNR adjusted)” (gray)indicates the projected variance after the correction for potential SNR difference between mouse and marmoset imaging. Because of the difference in FOV size, the number of pixels allocated to one neuron is larger for the mouse than for the marmosets, which could yield a better SNR for the mouse. To rule out the possibility that this difference affected the overall results, we re-analyzed the mouse data, with the number of pixels allocated to each cell being comparable to that of the marmoset. The median numbers of pixels allocated to each cell in the marmoset and mouse V1 were 110 and 208, respectively. To match the number of pixels allocated to each cell in mouse V1, we randomly selected 100 pixels within the original cell-mask and used them to obtain new cell time courses. These new cell time courses were then used to calculate the stimulus-related variance projected onto a shared space. The overall results of this control analysis did not differ from the original results; the value of the projected variance obtained with the new time courses of the mouse V1 was smaller than that of the marmoset visual cortex. Moreover, the projected variance of the mouse V1 calculated using the new and original time courses did not show a statistically significant difference (*p* > 0.1, Wilcoxon signed-rank test).

**Extended Data Figure 12** | **Estimated numbers of dimension of shared space and control analysis with fixed number of trials and cells. a**) Numbers of dimensions of shared space across visual areas. **b-c**) Control analysis conducted with fixed number of trials (20) and cells (400). (b) shows factions of stimulus related variance projected onto shared spaces in marmoset V1 and mouse V1. The amount of the projected variance was significantly larger in marmoset V1 than in mouse V1 (p < 0.00001, two-sample t test). (c) Estimated numbers of dimensions of shared space in marmoset V1 and mouse V1. Estimated numbers of dimensions were significantly larger in marmoset V1 than in mouse V1 (p< 0.03, two-sample t test).

**Extended Data Figure 13** | **Examples of spontaneous PCs in widefield imaging**. Top 20 PCs of spontaneous activity are shown for two example FOVs. **a**) V1/V2 FOV. **b**) MT FOV.

**Extended Data Figure 14** | **Stimulus-related variance projected onto 100-dimensional shared space**. Left, same convention as Figure 4a but projected variances were calculated using 100-dimensional shared space. Each point indicates FOV. Right, same as the left but the data are summarized by animals.

**Extended Data Figure 15** | **A neural network simulation that tests effect of structured versus examined functional architecture on the orthogonality between spontaneous and evoked activity. a**) Model architectures. Two-layer fully connected networks are used. Each unit has preferred orientation. Units in the marmoset V1 model (left) are spatially structure by preferred orientation, mimicking columnar structure. Units in the mouse V1 model (right) are spatially disorganized, mimicking salt-and-pepper organization. Inset in the bottom left shows examples of inputs to L1. In the simulation of spontaneous activities, L1 units received structured noise inputs. In the simulation of visually evoked activities, L1 units received orientation-dependent inputs plus structured noise inputs. Note that structured noise inputs are added to simulate single-trial activity. Connection weights between L1 and L2 units depend on difference of the preferred orientation between units (inset in the right bottom). Weight profile of marmoset V1 is made sharper than that of mouse V1 to resemble structured organization in marmoset V1. **b)** Example snapshots of spontaneous activity in L2 (right panels). Unit-wise correlations of spontaneous activity are also shown (left panels). **c)** Same as (b) but for visually evoked activities. **d)** Orientation tuning curves in example L2 units. **e)** Unit-wise correlation between spontaneous activities and trial-averaged visually evoked activities in L2. Same analysis and figure convention as in Fig. 2g. **f)** Fraction of stimulus related variance projected to shared spaces. Same analysis as in Fig. 4a. Histogram of the fractions across 100 simulations are shown.

## Supplementary Video Legends

**Supplementary Video 1** | **Spontaneous cortical activity in occipital and parietal areas**. Movie of spontaneous cortical activity in FOV shown in **Fig. 1b-d**. Note that no high-pass filtering is used.

**Supplementary Video 2** | **Spontaneous cortical activity in V1 and V2**. Movie of spontaneous cortical activity in FOV shown in **Extended Data Fig. 2d-f**. Note that no high-pass filtering is used. Note that no high-pass filtering is used.

**Supplementary Video 3** | **Spontaneous cortical activity in temporal areas**. Movie of spontaneous cortical activity in FOV shown in **Fig. 1e-g**. Note that no high-pass filtering is used. Note that no high-pass filtering is used.

**Supplementary Video 4** | **Another example of spontaneous cortical activity in occipital and parietal areas**. Movie of spontaneous cortical activity (not filtered) in FOV shown in **Extended Data Fig. 1b-d**. Note GCaMP6f was used in this example.

