## Supporting Discussion for "Orthogonalization of spontaneous and stimulus-driven activity by hierarchical neocortical areal network in primates"

**Supporting Discussions**

**Supporting Discussion 1**

In the marmoset V1, spontaneous and visually evoked activity patterns were similar. On the other hand, in the mouse V1, the two types of activity patterns were dissimilar. This species difference may be related to another species difference of V1: the presence and absence of orientation columns in marmosets and mice, respectively (*1-3*). Neurons sharing the same preferred orientation tend to have high noise correlation (*4, 5*) as well as spontaneous correlation (*6*). In the marmoset V1 in which orientation columns are present, local synaptic inputs to a single neuron are likely to share similar preferred orientation (*7, 8*). Hence, a marmoset V1 neuron tends to receive correlated spontaneous synaptic inputs. The correlated activity of presynaptic neurons activates the postsynaptic neuron sharing the same preferred orientation. Such cascade of activity likely explains correlated spontaneous activity of neurons sharing similar preferred orientation. In contrast, because of the absence of orientation columns, a mouse V1 neuron tends to receive local inputs from neurons whose preferred orientation are diverse. Thus, a mouse V1 neuron likely receives less correlated spontaneous synaptic inputs compared with a marmoset V1.

To test the validity of this hypothesis, we created simple two-layer neural network models that mimicked the marmoset or mouse V1 and examined whether a more disordered structure in mice leads to the orthogonalization of spontaneous and stimulus-driven activity (Extended Data Figure 15)(see section “Details of Neural Network Simulations” below for details). A two-layer fully connected network was used as the base architecture for both the marmoset and the mouse V1 models. In this model, the visual inputs were orientation-selective neural activity (such as the neural activity in layer 4 of mouse V1) (L1 in Extended Data Figure 15a). The neurons in the input layer are connected to the top layer (L2) in an all-to-all fashion but with different weights. Ordered vs. disordered architectures (i.e. columnar vs. salt-and-pepper architectures) of the marmoset and mouse V1, respectively, are modeled by two distinct profiles of connection weights as functions of difference in preferred orientations of the connected neurons (right panels in Extended Data Figure 15a). The marmoset V1 model has a narrowly tuned weight profile with respect to difference in preferred orientations of the connected neurons, so that a neuron in L2 preferentially samples inputs from L1 neurons sharing similar preferred orientation. In contrast, the mouse V1 model has a broadly tuned weight profile, such that a neuron in L2 samples inputs from L1 neurons with diverse preferred orientations. The weight profile of the mouse V1 model was chosen based on previous physiological studies reporting that mouse L2/3 neurons receive broadly orientation selective inputs (*4*).

We ran simulations of visually evoked activity and spontaneous activity using these networks (bottom panels in Extended Data Figure 15a): For the simulation of spontaneous activity, we injected a structured noise to L1 neurons. The structured noise is modeled based on previous reports showing similarity of signal correlation and noise correlation. In both primates and rodents, previous studies showed that V1 neurons with high signal correlation, hence similar preferred orientations, also showed high noise correlation and spontaneous activity correlation (Kohn & Smith, J. Neurosci., 2005; Ko et al., Nature, 2011). To simulate these previous observations, we convolved a white noise by using a Von Mises function to introduce noise correlation between L1 units sharing similar preferred orientations. Extended Data Figure 15b shows examples of simulated spontaneous activity in the mouse V1 and marmoset V1 models. Although the structured noise inputs to L1 neurons are the same for the marmoset V1 model and the mouse V1 model, correlated activities in L1 neurons are weighted more by each L2 neuron in the marmoset V1 model than the mouse V1 model because of a narrower tuning of the weight profile in the marmoset V1 model. Thus, L2 neurons sharing similar preferred orientations in the marmoset V1 model should show correlated spontaneous activity. On the other hand, L2 neurons sharing similar preferred orientation in the mouse V1 model should show less correlated spontaneous activities, because each L2 neuron put more weights on diverse (less correlated) inputs from L1 neurons. Consistently, the activity correlation between L2 neurons revealed a more structured correlation matrix for the marmoset V1 model than for the mouse V1 model (right panels in the new Extended Data Figure 15b). To simulate visually evoked activity, in addition to structured noise, we injected activity tuned at a selected preferred orientation into L1 neurons, mimicking the presentation of oriented gratings. Extended Data Figure 15c shows examples of simulated visually evoked activity in the mouse V1 and marmoset V1 models. Activity correlation between L2 neurons (“signal correlation”) was similar for the two models (right panels in Extended Data Figure 15c). The orientation tuning curve of each L2 neuron was narrower in the marmoset V1 model than in the mouse V1 model, mimicking previous animal studies (*9, 10*). These correlation structures suggest that spontaneous activity patterns and visually evoked activity patterns were similar, at least in the correlation structure, in the marmoset V1 model but not in the mouse V1 model.

To confirm that the simulation reproduced essential patterns of the animal results, using the simulated data, we conducted the same analyses that we used for the animal data. As shown in the Figure 2g, the maximum correlation value between the spontaneous and evoked-activity frames were significantly higher in the marmoset V1 model than in the mouse V1 model (*p* < 10^-42^, rank-sum test; Extended Data Figure 15e). Similarly, as in shown in Figure 4a, the fraction of the variance of visually evoked activity projected to the shared space was larger for the marmoset V1 model than for the mouse V1 model (*p* < 10^-33^, rank-sum test across 100 instances of model pairs; Extended Data Figure 15f). These simulation results are consistent with animal data and suggest that spontaneous and visually evoked activity are more orthogonalized in the mouse V1 model than in the marmoset V1 model. Taken together, these results suggest that a more disordered connection leads to greater orthogonalization between spontaneous and evoked activities.

*Details of Neural Network Simulations*

Model architectures

Neural network simulations were conducted using Matlab (MathWorks, Natick, MA). The number of neurons in each model layer was set to 180. For each neuron, we assigned a unique preferred orientation $\theta$ ranging from 1 to 180 degrees.

The connection weight (W) between the $\theta$-th neuron in layer 2, whose preferred orientation is $\theta$ deg, and a layer 1 neuron, whose preferred orientation is $\varphi$ deg, was determined by a normalized Von Mises function as follows:

$${W'}_{x}=exp\left( A*\left( \cos\left( \frac{x*pi}{90} \right)-1 \right) \right) (1)$$

$$W_{\theta-\varphi}=\frac{{W'}_{\theta-\varphi}}{\sum_{i} {W'}_{i}} (2)$$

where A is a parameter of the Von Mises function, and an index i in the summation spans from 1 to 180 at a step equal to 1.

We separately set the connection weight W for the marmoset (W_marmoset_) and the mouse (W_mouse_) models, such that the mouse layer 2 neurons receive broader orientation-selective inputs than the marmoset model; A = 1.8 for W_marmoset_; A = 0.6 for W_mouse_ (see Extended Data Fig. 15a).

Generation of visually-evoked and spontaneous activities

For the generation of spontaneous activity, we first generated random noise activities independently sampled from a normal distribution:

$$\mathbf{n}_{0}=\left\{ n_{0\_1},n_{0\_2},\ldots,n_{0\_\theta} \right\}, n_{0\_\theta}\mathcal{\sim N}\left( 0,1 \right) (3)$$

These activities were rectified by a threshold value t_0_ (t_0_ = 0.1):

$$\mathbf{r}_{0}=relu\left( \mathbf{n}_{0}\boldsymbol{,}t_{0} \right) (4)$$

Then, these activities were convolved with a weight kernel w:

$${w'}_{x}=exp\left( \cos\left( \frac{x*pi}{90} \right)-1 \right) (5)$$

$$w_{x}=\frac{{w'}_{x}}{\sum_{i} {w'}_{i}}*0.5 (6)$$

$$r_{0\_\theta}=\sum_{k=\theta-90}^{\theta+89} w_{mod(k,180)}*r_{0\_k} (7)$$

where an index i runs from 1 to 180 at a step equal to 1.

Finally, a random noise was added to the each of the activity with an arbitrary coefficient to obtain input to layer 1 neurons ($\mathbf{r}_{\mathrm{sponta}}$):

$$\mathbf{n}_{1}=\left\{ n_{1_{1}},n_{1_{2}},\ldots,n_{1_{\theta}} \right\}, n_{1_{\theta}}\mathcal{\sim N}\left( 0,1 \right) (8)$$

$$\mathbf{r}_{\mathrm{sponta}}=\mathbf{r}_{0}\boldsymbol{+}\mathbf{0.05}\boldsymbol{*}\mathbf{n}_{1} (9)$$

For the generation of visually-evoked activity, we first generated a spontaneous input to L1 neurons ($\mathbf{r}_{\mathrm{sponta}}$), created as described above, and then generated a visual input ${\mathbf{V}=\{V}_{1}, V_{2},\ldots,V_{180}\}$ as explained below.

Visual input with the stimulus orientation $\vartheta$ deg to the $\theta$-th neuron in layer 2, whose preferred orientation is $\theta$ deg, was defined by the Von Mises function:

$${V'}_{\theta,\vartheta}=exp\left( 2.5* \left( \cos\left( \frac{\left( \theta-\vartheta\right)*pi}{90} \right)-1 \right) \right) (10)$$

$$V_{\theta,\vartheta}=\frac{{V'}_{\theta,\vartheta}}{\sum_{k} V_{k,\vartheta}} (11)$$

where an index k runs from 1 to 180 at a step equal to 1. The stimulus orientations ($\vartheta$) spanned from 1 deg to 180 deg at a 1 deg step (i.e. $\vartheta\in\{1,2,\ldots,180$}).

Then, a visual input L1 neurons ($\mathbf{r}_{\mathbf{vis}}$) was obtained using a following equation:

$$\mathbf{r}_{\mathbf{vis}}=\mathbf{r}_{\mathrm{sponta}}+9.0*\mathbf{V} (12)$$

Activities of layer 1 neurons (**R**_1_) were obtained by rectifying inputs to L1 neurons by a threshold value t_1_ (t_1_ = 0.1):

$$\mathbf{R}_{1}=relu\left( \mathbf{r}_{\mathrm{input}}\boldsymbol{,}t_{1} \right) (13)$$

$$\mathrm{where}\mathbf{r}_{\mathrm{input}} \mathrm{is}\mathbf{r}_{\mathbf{sponta}}\mathrm{and}\mathbf{r}_{\mathbf{vis}} for spontaneous and visually evoked activities, respectively.$$

An input from layer 1 neurons to the $\theta$-th neuron in layer 2, whose preferred orientation is $\theta$ deg, can be expressed as:

$${r2}_{\theta}=\sum_{k=\theta-90}^{\theta+89} W_{k}*{R1}_{k} (14)$$

where the ${\mathbf{R1}=\{R1}_{1}, {R1}_{2},\ldots,{R1}_{180}\}$ and the ${\mathbf{r2}=\{r2}_{1}, {r2}_{2},\ldots,{r2}_{180}\}$ correspond to the activity (output) of layers 1 neurons and inputs to 2 neurons, respectively.

Next, a random noise was further added to the layer 2 neuron input with:

$$\mathbf{r2}_{\mathrm{noisy}}= \mathbf{r2}+0.05 *\mathbf{n}_{2} (15)$$

$$\mathbf{n}_{2}=\left\{ n_{2\_1},n_{2\_2},\ldots,n_{2\_\theta} \right\}, n_{2\_\theta}\mathcal{\sim N}\left( 0,1 \right) (16)$$

Finally, the layer 2 neuron activity was rectified by a threshold value t_2_ (t_2_ = 0.1):

$$\mathbf{R2}=relu\left( \mathbf{r2}_{\mathrm{noisy}}\boldsymbol{,}t_{2} \right) (17)$$

For visually evoked activity, total of 1000 visual stimulation runs, yielding 180,000 activity patterns (which we call frames) were generated. For spontaneous activity, total of 1000 frames of spontaneous activity simulation were generated. In both visual stimulation runs and spontaneous activity runs, individual frames were treated as independent. A code used for the simulation is available for download (https://github.com/teppei-matsui/NN_Simulation).

Orientation tuning width for each L2 unit was obtained by fitting Von Mises function to trial averaged orientation responses, with A, W and $\theta$ as free parameters:

$$VM(x,A,W,\theta)= A* exp \left( W \left( \cos\left( \frac{\left( x-\theta\right)*pi}{90} \right)-1 \right) \right)$$

Analysis of simulated spontaneous and visually evoked activities were conducted similarly as for mouse and marmoset data. In the calculation of the fraction of visual stimulus-related variance projected to the shared space (Extended Data Fig. 15f), we used 20 spontaneous PCs to obtain shared space. 100 instances of simulations were conducted to generate distribution of the fraction of projected variances. See simulation code attached below for further details.

**Supporting Discussion 2**

The progressive orthogonalization of spontaneous and stimulus evoked activity patterns may be explained by the concept of communication subspace of cortico-cortical connections (*11, 12*). In the macaque visual cortex, V1 activity patterns contained in the communication subspace are more likely to evoke downstream activity in V2 (*11*). We conjecture that the activity patterns of stim-only PCs are more effective in activating neurons in the downstream area, thus constitute the communication subspace. In contrast, the activity patterns of shared PCs are less effective in activating down-stream neurons. After multiple steps of cortico-cortical activity propagations, activity patterns corresponding to shared PCs are diminished whereas the patterns corresponding to stim-only PCs are efficiently propagated.
